# Exploring the niche concept in a simple metaorganism

**DOI:** 10.1101/814798

**Authors:** Peter Deines, Katrin Hammerschmidt, Thomas CG Bosch

## Abstract

Organisms and their resident microbial communities - the microbiome - form a complex and mostly stable ecosystem. It is known that the composition of the microbiome and bacterial species abundances can have a major impact on host health and *Darwinian fitness*, but the processes that lead to these microbial patterns have not yet been identified. We here apply the niche concept and trait-based approaches as a first step in understanding the patterns underlying microbial community assembly and structure in the simple metaorganism *Hydra*. We find that the carrying capacities in single associations do not reflect microbiota densities as part of the community, indicating a discrepancy between the fundamental and realized niche. Whereas in most cases, the realized niche is smaller than the fundamental one, as predicted by theory, the opposite is observed for *Hydra*’s two main bacterial colonizers. Both, *Curvibacter* sp. and *Duganella* sp. benefit from association with the other members of the microbiome and reach higher fractions as compared to when they are the only colonizer. This cannot be linked to any particular trait that is relevant for interacting with the host or by the utilization of specific nutrients but is most likely determined by metabolic interactions between the individual microbiome members.

## Introduction

Microbiomes contribute to ecosystems as key engines that power system-level processes (Falkowski et al., 2008). This also applies to host ecosystems, where they are critical in maintaining host health, survival, and function (Kau et al., 2011; McFall-Ngai et al., 2013). Despite their importance, the mechanisms governing microbiome assembly and composition are largely unknown. This is different for macroscopic communities, thanks to the application of niche (Holt, 2009; Leibold, 1995; Whittaker et al., 1973) and trait-based theories, which might also provide a useful framework for studying the ecology and evolution of microbiomes in metaorganisms (Kopac and Klassen, 2016).

The niche concept is one of the core concepts in ecology and has been rediscovered by modern ecology for explaining biodiversity and species coexistence patterns (Pocheville, 2015). The niche-based theory states that an ecological community is made up of a limited number of niches, each occupied by a single species. Hutchinson (Hutchinson, 1957) defined the *fundamental niche* as the needs of a species for it to maintain a positive population growth rate, disregarding biotic interactions (Hutchinson, 1957; Pearman et al., 2008). The fundamental niche therefore represents an idealized situation exclusive of interspecific interactions. The effect of biological interactions is taken into account in the definition of the *realized niche* (Hutchinson, 1957). This is the portion of the fundamental niche in which a species has a positive population growth rate, despite the constraining effects of biological interactions, such as inter-specific competition (Hutchinson, 1957; Pearman et al., 2008).

In the last two decades, the shift from taxonomy to function by using trait-based approaches has provided a detailed understanding of biodiversity-ecosystem functioning (Louca et al., 2018). Recently, this framework is also being used by microbial ecologists to study microbial biogeography (Green et al., 2008), or to unravel microbial biodiversity-ecosystem functioning relationships (Krause et al., 2014). Further, this approach allows studying microbiomes in the light of coexisting traits/ functions rather than of coexisting microbes (Martiny et al., 2015). A recent study successfully used this approach and analyzed trait-based patterns to understand the mechanisms of community assembly and succession of the infant gut microbiome (Guittar et al., 2019). Microbial traits cover a range of phenotypic characteristics, for example organic phosphate utilization, bacteriophage host range, cellulose degradation, biofilm formation, nitrogen fixation, methanogenesis, and salinity preference (Martiny et al., 2015). Potential microbial traits can be measured directly by laboratory assays (as in this study) or can be indirectly inferred based on genomic information.

The aim of this study is to apply the niche concept and trait-based theory to the metaorganism *Hydra vulgaris* (strain AEP) to gain insight into the mechanisms underlying the microbial community composition. We thus specifically extend the niche-assembly perspective, classically used for assessing species assembly and coexistence in abiotic environments, to a host-associated microbiome, thus a biotic environment.

The freshwater polyp *Hydra* and its microbiome have become a valuable model system for metaorganism research as it provides a bridge between the simplicity of synthetic communities and the complex mouse model (Deines and Bosch, 2016). The ectoderm is covered by a multi-layered glycocalyx, which is the habitat for a highly stable, low complexity, species specific microbiome (Bosch, 2013; Deines et al., 2017; Franzenburg et al., 2013), with a carrying capacity of 1.7*10^5^ CFUs per *Hydra* (Deines et al., 2020). The six most abundant bacterial colonizers of *Hydra vulgaris* (strain AEP) make up between 84 - 90% of *Hydra*’s microbiome (Franzenburg et al., 2013; Murillo-Rincón et al., 2017). These species, *Curvibacter* sp. (65 - 76%), *Duganella* sp. (11 - 16%), *Undibacterium* sp. (1 - 2%), *Acidovorax* sp. (0.4 - 0.7%), *Pseudomonas* sp. (0.4%), and *Pelomonas* sp. (0.2 - 0.9%), were isolated and purified by Fraune et al. (2015). The single-species isolates can be cultured and manipulated *in vitro* (Bosch, 2013; Fraune et al., 2015; Wein et al., 2018), allowing the measurement of phenotypic microbial traits and fitness. Fitness, as defined by niche theory, is the positive population growth of the focal species, which in our study is that of the six available microbiome members. Measurements of the performance of the bacterial populations when grown singly, i.e. in the absence of the other microbial competitors *in vitro* and *in vivo* (on germ-free *Hydra* polyps), specify the fundamental niche. For each species, we compare the fundamental niche to the realized niche, which we calculated based on published data on the microbiome composition of wild-type and conventionalized polyps (germ-free polyps incubated with tissue homogenates of wild-type animals) (Franzenburg et al., 2013; Murillo-Rincón et al., 2017). We also measure phenotypic traits that might be connected to the success of the various microbial species in occupying the fundamental niche; these are essentially traits that might play a role in successfully populating their environment, the host, such as biofilm formation, surface hydrophobicity (bacterial cells are more likely to attach to surfaces with the same hydrophobicity), and nutrient utilization patterns. Here we focused on carbon sources as the microbiome inhabits the outer mucus-like layer of *Hydra*’s glycocalyx (Fraune et al., 2015), which is carbohydrate-rich (Ouwerkerk et al., 2013; Schröder and Bosch, 2016). As the realized niche is determined by biological interactions of one species with its associate microbial community, we focus on traits that are important when competing with other species, such as growth rate, niche overlap, and niche breadth. Ultimately, we take the traits and the ecological niches as determinants of species interactions, which may infer the assembly and structure of the host-associated microbiome.

## Materials and Methods

### Animals used, culture conditions, and generation of germ-free animals

*Hydra vulgaris* (strain AEP) was used in the experiments and cultured according to standard procedures at 18°C in standardized *Hydra* culture medium (Lenhoff and Brown, 1970). Animals were fed three times a week with 1st instar larvae of *Artemia salina*. Germ-free polyps were obtained as previously described (Franzenburg et al., 2013; Murillo-Rincón et al., 2017). After two weeks of treatment, polyps were transferred into antibiotic-free sterile *Hydra* culture medium for recovery (four days). Sterility was confirmed by established methods (Franzenburg et al., 2013). During antibiotic treatment and re-colonization experiments, polyps were not fed.

### Bacterial species and media

The bacterial species used in this study are *Curvibacter* sp. AEP1.3, *Duganella* sp. C1.2, *Undibacterium* sp. C1.1, *Acidovorax* sp. AEP1.4, *Pelomonas* sp. AEP2.2, and *Pseudomonas* sp. C2.2 (all Gammaproteobacteria), all of which were isolated from the *Hydra vulgaris* (strain AEP) microbiome (Fraune et al., 2015). These bacteria were cultured from existing isolate stocks in R2A medium at 18°C, shaken at 250 r.p.m for 72 h before use in the different experiments.

### Fundamental and realized niche

Germ-free polyps (n=18) were inoculated with single bacterial species using 5×10^3^ cells in 1.5 ml Eppendorf tubes containing 1 ml of sterile *Hydra* culture medium (each n=3). After 24 h of incubation, all polyps were washed with, and transferred to sterile *Hydra* culture medium, incubated at 18°C. After three days of incubation individual polyps were homogenized in an Eppendorf tube using a sterile pestle, after which serial dilutions of the homogenate were plated on R2A agar plates to determine colony-forming units (CFUs) per polyp (Deines et al., 2020). The dilution with counts in the countable range was selected for each bacterial species, counting a minimum of 50 CFUs per plate.

The carrying capacities of mono-associations provide information of the occupied niche space on the host in the absence of other microbial species that are part of the microbiome, and thus specifies the fundamental niche for each (as calculated from the proportion of each species from the sum of all).

As microbial community composition and relative microbial abundances of wild-type and conventionalized polyps have been reported to be remarkably stable over time (Bosch, 2013; Franzenburg et al., 2013; Fraune et al., 2015; Murillo-Rincón et al., 2017), we here base the estimates of the realized niche on the underlying original data from previous studies (Franzenburg et al., 2013; Murillo-Rincón et al., 2017). For OTU (operational taxonomic unit) estimations of *Hydra*’s microbiome, both studies used amplicon sequencing of the variable regions 1 and 2 (V1V2) of the bacterial 16S rRNA genes (V1V2-one step approach). The sequences were grouped into OTUs at a ≥ 97% sequence identity threshold. For the relative abundance estimates, all samples were normalized to the lowest number of reads in each respective dataset. Note that it has been shown for *Hydra*’s microbiome that relative abundance estimations based on 16S rRNA amplicon sequencing do not significantly differ compared to relative abundance estimations based on metagenomic shotgun sequencing (Rausch et al., 2019).

Carrying capacities of the members when part of the full community could not be assessed via plating of the community as was done for the mono-associations as not all bacterial species can be differentiated based on colony morphology, which is why we based calculations of the realized niche on data that was previously published (Franzenburg et al., 2013; Fraune et al., 2015; Murillo-Rincón et al., 2017). Other studies demonstrate that germ-free polyps, which were inoculated with the natural community of species (conventionalized animals), harbor an equally dense microbiome (CFUs per polyp; Deines et al., 2020) that is remarkably similar in its composition (Murillo-Rincón et al., 2017) as compared to wild-type polyps. This indicates that the generation and re-exposure of germ-free animals does not lead to an overall change in carrying capacity or community composition.

### Cell surface hydrophobicity (CSH)

The BATH assay was performed as described previously (Borecká-Melkusová and Bujdáková, 2008; Rosenberg, 1984). It uses a biphasic separation method to measure cell surface hydrophobicity. In short, for each species tested, exponential growth phase cultures of the six species were adjusted to an optical density at 600nm (OD_600_) of 0.1 in R2A medium (OD_initial_). 4 ml of this bacterial suspension was placed into a glass tube, overlaid with 1 ml of n-hexodecane (Sigma Aldrich), and vortexed for 3 min. The phases were then allowed to separate for 15 min, after which the ODs of the aqueous (lower) phase containing hydrophilic cells was measured at OD_600_ (OD_residual_). The hydrophobic cells are found in the n-hexodecane overlay (upper phase). OD values were compared to the bacterial suspension before mixing with n-hexodecane. The relative hydrophobicity (RH) was calculated as follows: RH = ((OD_initial_ – OD_residual_)/OD_initial_) x 100%. The experiment was performed in triplicate with independent bacterial overnight cultures.

### Biofilm quantification by use of crystal violet (CV)

Biofilm formation was assayed and quantified as previously described (Ren et al., 2015). Briefly, exponential growth phase cultures of the six species were adjusted to an OD_600_ = 0.1 in R2A medium. Biofilm formation was assayed in a 96 well plate using four replicates for each treatment inoculated from the same bacterial overnight culture. For single isolates an inoculation volume of 180 µl was used. After 48 h of incubation at 18°C with shaking (200 r.p.m) biofilm formation was quantified by a modified crystal violet (CV) assay (Peeters et al., 2008).

### Characterizing nutrient (carbon) utilization

To characterize the nutrient profiles, specifically the carbon metabolism profile, for each species of *Hydra*’s microbiota, we used BIOLOG GN2 plates. BIOLOG GN2 plates are 96-well microwell plates containing 95 different carbon sources plus a carbon-absent water control well. Species were grown from isolate stocks in R2A medium (18°C, shaken at 250 r.p.m.), centrifuged at 3000 r.c.f. for 5 min, re-suspended in S medium and adjusted to an OD_600_ of 0.1. Each well of the BIOLOG plate was inoculated with 150 µl of bacterial suspension and incubated for three days at 18°C in a humid chamber. Growth on each of the 95 nutrients was determined as OD_600_ of each well using a TECAN plate reader. For each plate, the OD of the water control was subtracted from the reading of all other wells prior to analysis, and differenced OD values below 0.005 were considered as no growth (Vaz Jauri et al., 2013). Nutrient use was evaluated on three replicate plates inoculated from independent bacterial overnight cultures. Nutrient niche overlap (NO) was calculated using the formula: NO = (number of nutrients used by both A and B)/((number of nutrients used by A + number of nutrients used by B)/ 2) (Vaz Jauri et al., 2013). A value of 1 indicates the use of the same nutrients (100% overlap) and 0 indicates no nutrient overlap among the 95 substrates tested. We also calculated the relative use of the eleven functional groups (carbohydrates, carboxylic acids, amino acids, polymers, aromatic chemicals, amines, amides, phosphorylated chemicals, esters, alcohols and bromide chemicals) according to Daou et al. (2017). In brief, the relative use of C substrates was calculated as absorption values in each well divided by the total absorption in the plate.

### Measurement of bacterial growth rates *in vitro*

Cultures of the all six species were produced in R2A microcosms (grown for 72 h at 18°C, at 250 r.p.m). Aliquots of each culture were first washed in S medium and then re-suspended in fresh R2A medium to an optical density of 0.025 at 600 nm (OD_600_). Growth kinetics of all species were determined in 96-well microtiter plates. A 100 µl aliquot of each re-suspension was pipetted into 100 µl of fresh R2A medium. The microtiter plate was then placed in a microplate reader (TECAN Spark 10M, Tecan Group Ltd., Switzerland), and the OD_600_ of each well was measured at 30 min intervals for 96 cycles (with 10 sec shaking at 150 r.p.m. prior to each read). The growth of each species was determined in five well locations on an individual 96-well plate, which was replicated six times with independent bacterial overnight cultures. The maximum growth rate (µ_max_) was calculated from the maximum slope of the absorbance over time.

### Statistical analysis

Analysis of variance (ANOVA) and subsequent post hoc Tukey-Kramer tests were used to test for differences in the carrying capacity of the six *Hydra* colonizers. To meet the requirements for the model, the variable was Box-Cox transformed. A Welch ANOVA (and subsequent Wilcoxon post hoc tests) was used to test for differences in the fraction of the different species in the community, differences in biofilm formation capacity, and *in vitro* growth rates between the species. Analysis of variance (ANOVA) and subsequent post hoc Tukey-Kramer tests were used to test for differences in the cell surface hydrophobicity of the six *Hydra* colonizers.

Sample size was chosen to maximise statistical power and ensure sufficient replication. Assumptions of the tests, that is, normality and equal distribution of variances, were visually evaluated (see Quinn and Keough, 2003, for rationale). Non-significant interactions were removed from the models. Effects were considered significant at the level of P < 0.05. All statistical analyses were performed with JMP 9. Graphs were produced with GraphPad Prism 5.0, and RStudio (RStudio Team, 2015).

## Results

### Fundamental and realized niche occupation of the different bacterial species

In mono-colonizations, the six bacterial species differ significantly in their carrying capacity on *Hydra* (Figure 1A; ANOVA: *F*_5,12_=12.696, P=0.0002). The most extreme cases are *Acidovorax* sp. that reaches the highest numbers with 2.6*10^5^ CFUs/polyp, and *Duganella* sp. the lowest with 1.7*10^4^ CFUs/polyp. Mono-colonization of *Acidovorax* sp. thus exceeds the carrying capacity of the wild-type microbiome of 1.7*10^5^ CFUs per *Hydra* (Deines et al., 2020). Based on the carrying capacity of the single species in mono-colonisations, we estimated the fundamental niche of each of the six species. The realized niche of the six bacterial species is calculated based on previously published data on the composition of the extremely stable microbial community. The species differed in their relative abundance as part of the microbial community (Figure 1B; Welch ANOVA: *F*_5,14_=86.722, P<0.0001). The most dominant species is *Curvibacter* sp. representing 65% of the microbial community, followed by *Duganella* sp. that reaches about 16%. All other four species reach only comparatively low fractions (around 1%), with *Acidovorax* sp. being the lowest. When comparing the fundamental to the realized niche (Figure 2), we find that the realized niche of *Curvibacter* sp. and *Duganella* sp. is larger than their fundamental niche. This is in contrast to all other species, where as expected by theory, competition with other microbes leads to a smaller realized than the fundamental niche.

**Figure 1.**
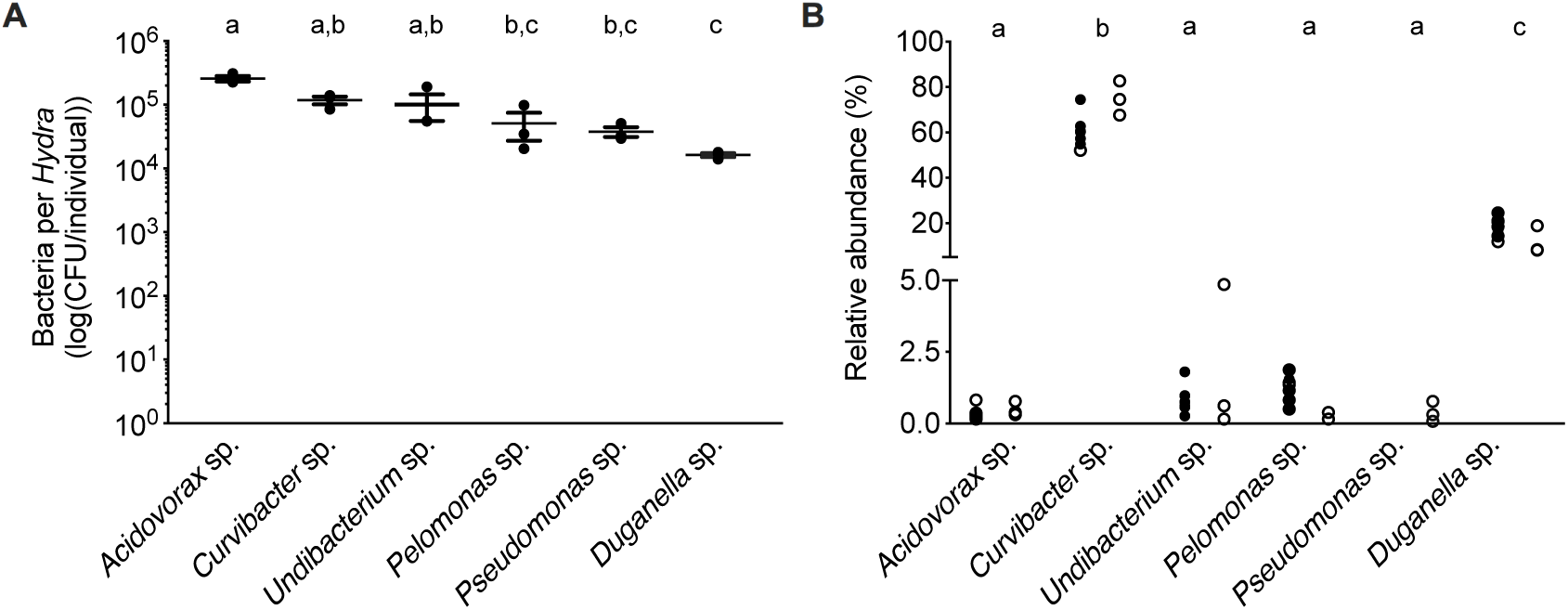
Performance of the six main bacterial colonizers isolated from the *Hydra* microbiome. (**A**) Carrying capacity of the *Hydra* ecosystem during mono-associations of germ-free polyps with individual bacterial species. Error bars are s.e.m., based on n=3. (**B**) Relative read abundances (%) of the different bacterial colonizers in wild-type (open circles) and conventionalized polyps (filled circles), compiled from previously published studies (left: (Murillo-Rincón et al., 2017), right: (Franzenburg et al., 2013)). Statistical differences as determined by post hoc tests (P < 0.05) are indicated by the different letters.

**Figure 2.**
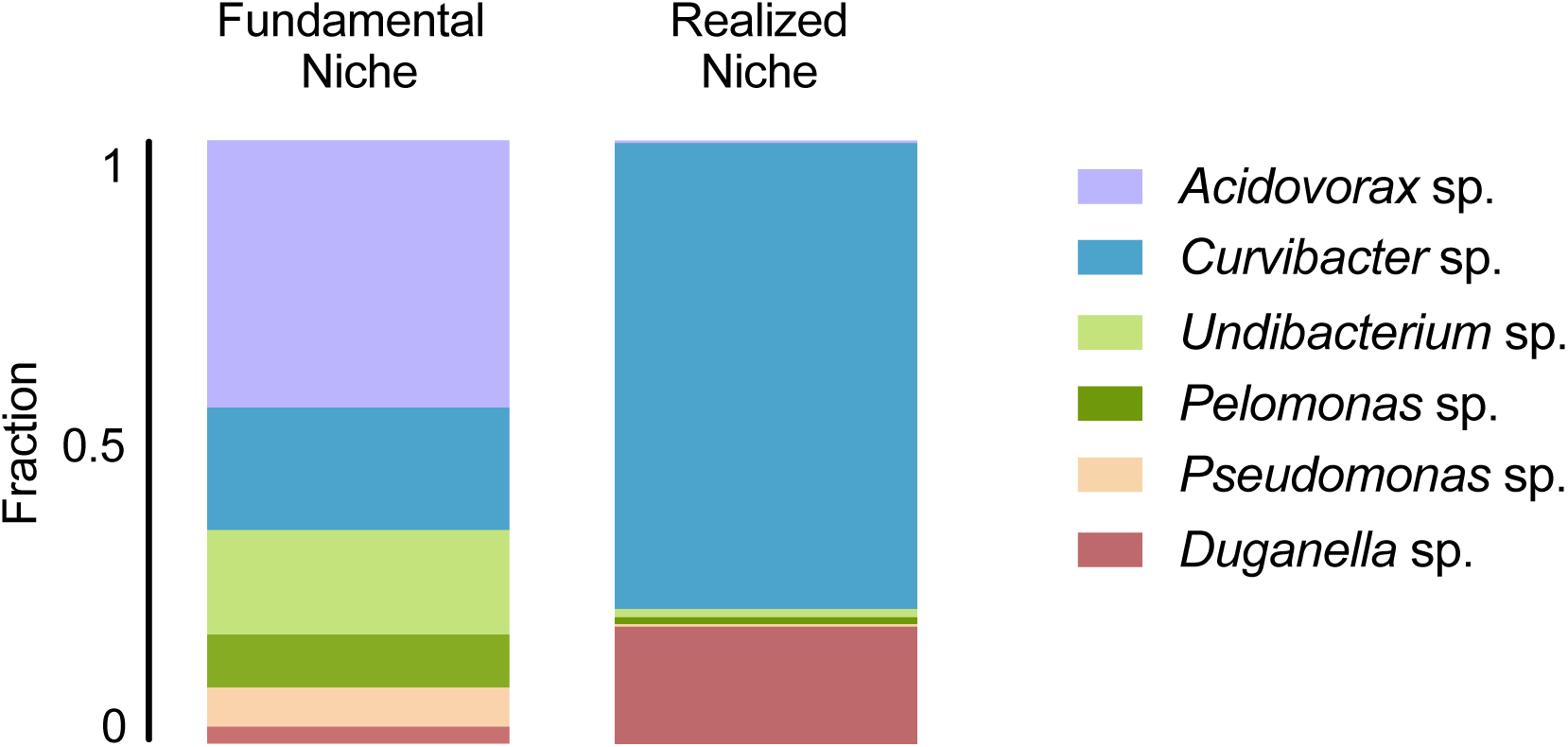
Fundamental and realized niches of six members of *Hydra*’s microbiome (based on mono-colonization’s and microbial community composition). The realized niche includes additional constrains arising from inter-specific competition between microbiome members.

### Bacterial traits

#### Associated with occupation of the fundamental niche

The BATH assay was conducted with six species of the *Hydra* microbiome to measure cell surface hydrophobicity (CSH). The CSH of the bacterial species vary, with values ranging from 0% to 42% and significantly differing between species (ANOVA: F_5,12_=26.869; P<0.0001). *Curvibacter* sp. and *Pelomonas* sp. were the only two species that did not show any affinity to the hexadecone; thus their cell population can be considered homogeneous consisting of only hydrophilic cells, which significantly differs from the others, except for *Pseudomonas* sp. (Figure 3A). *Pseudomonas* sp. and *Undibacterium* sp. show a mixed cell population, where 10 to 20% of the cells are hydrophobic. The species with the highest percentage of hydrophobic cells are *Acidovorax* sp. and *Duganella* sp., between 30 and 35%.

**Figure 3.**
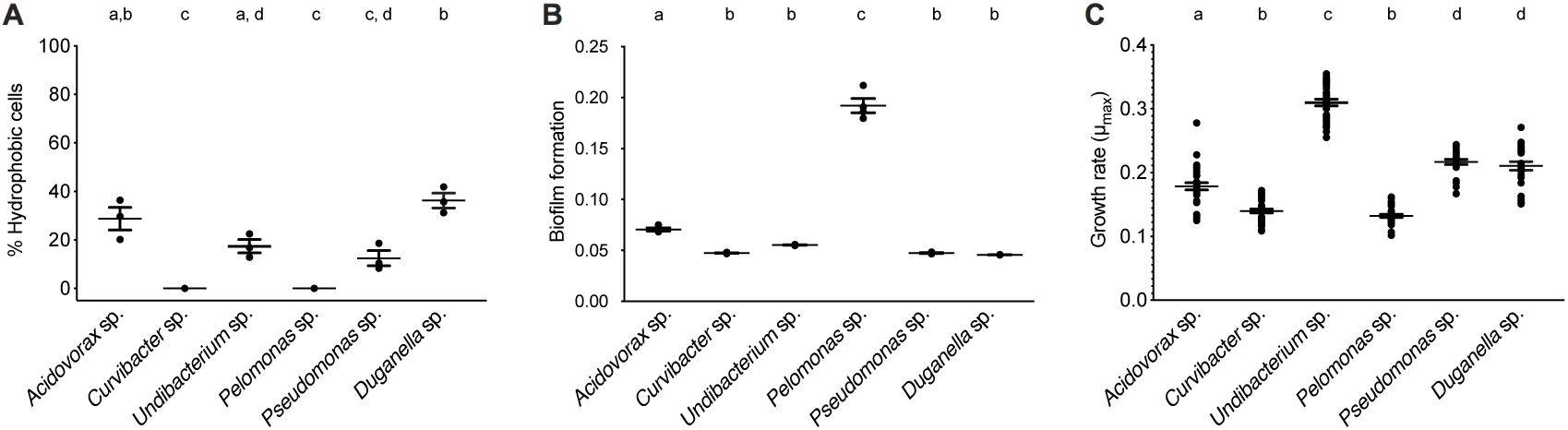
Trait measures of bacterial species isolated from the *Hydra* microbiome. (**A**) Cell surface hydrophobicity (Error bars are s.e.m., n=3), and (**B**) Biofilm formation capacity of six bacterial isolates. Error bars are s.e.m., n=4. (**C**) Bacterial growth rates of individual species measured *in vitro*. Error bars are s.e.m., based on n=30. Statistical differences as determined by post hoc tests (P < 0.05) are indicated by the different letters.

The species differed in their biofilm formation (Welch ANOVA: *F*_5,18_=350.723, P<0.0001). All species formed biofilms (Figure 3B), with *Pelomonas* sp. producing the largest biomass amount, which significantly differed from all other species. The biofilm amount of *Acidovorax* sp. was also significantly different from all other species but only roughly a third of the mass that *Pelomonas* sp. produced. All other species did not differ and are comparatively weak biofilm producers.

Nutrient utilization, i.e. carbon substrate usage, of all species was determined using a BIOLOG assay. The 94 carbon substrates are organized into eleven functional groups (Figure 4). Results showed that all six species actively oxidize carbon compounds such as carbohydrates (30-50% relative use), carboxylic acids (15-35% relative use) and amino acids (15-35% relative use) (Figure 4; Figure 5A). Carbohydrates are being used to an equal extent between all species, except for *Undibacterium* sp., which uses the highest amount of around 50% (relative use). Turanose is the compound most highly utilised, followed by a-D-lactose, L-rhamnose, and D-cellobiose. The use of carboxylic acids increases in the species, which are characterized by low frequencies in the *Hydra* microbiome, with the exception of *Duganella* sp. (with a relative use of 25-30%). Here D-galactonic acid lactone is the substrate with the highest usage, followed by different forms of hydroxyl butyric acids. Amino acids are most excessively used by *Acidovorax* sp., *Curvibacter* sp. and *Pseudomonas* sp., whereas the other species use amino acids to a lesser extent. Polymers are being used very differently between the species with *Pseudomonas* sp. showing the highest and *Undibacterium* sp. the lowest values. Amines are being used more frequent by the dominant species in the microbiome and are utilized to a lesser extent by the low abundant species.

**Figure 4.**
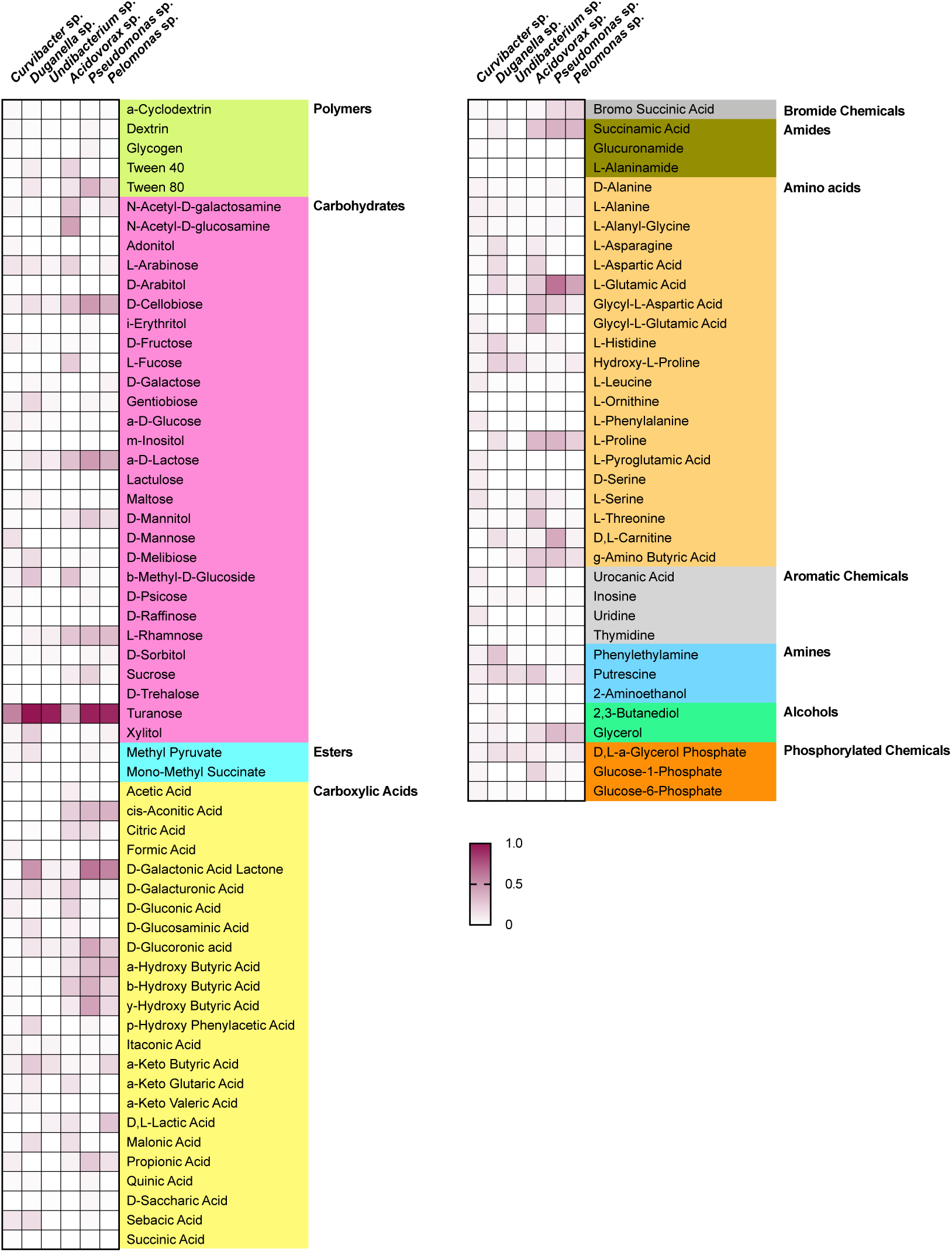
Substrate utilization pattern of six bacterial isolates from the *Hydra* microbiome measured with a BIOLOG assay. Colors indicate the relative magnitude of substrate utilization.

**Figure 5.**
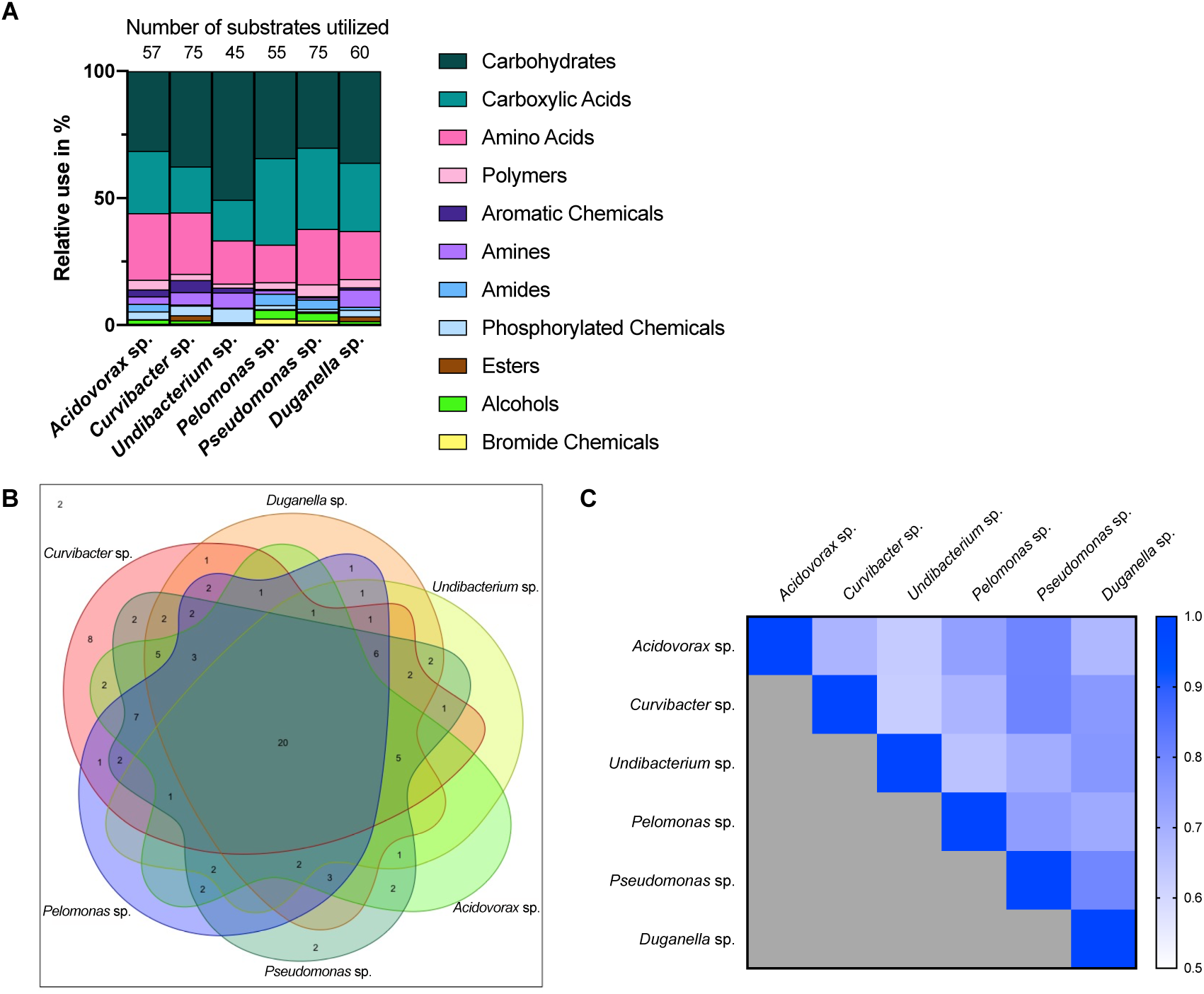
Metabolic phenotype diversity in *Hydra*’s microbiome. (**A**) Number of substrates utilized (out of 95) and their relative use (%). (**B**) Venn diagram showing the distribution of shared substrates among the microbiome members. (**C**) Niche overlap among all pairwise combinations of six *Hydra* microbiome members. A value of 1 indicates the use of the same nutrients (100% overlap) and 0 indicates no nutrient overlap.

#### Associated with occupation of the realized niche

When comparing the *in vitro* growth rates we find species perform differently (Figure 3C; Welch ANOVA: *F*_5,174_=223.856, P<0.0001). The fastest species, *Undibacterium* sp., grows twice as fast as compared to the slowest one, *Pelomonas* sp..

The overlap in carbon substrate usage between all six microbiome members is displayed as a Venn diagram (Figure 5B). There are only two substrates, which are not utilized by any species, whereas 20 substrates are used by all species. There are only two species that can metabolize substrates that none of the other species is using. While *Pseudomonas* sp. uses two substrates: i-erythritol and lactulose, *Curvibacter* sp. is able to utilise eight substrates: D-arabitol, D-mannose, D-trehalose, mono-methyl succinate, formic acid, glucuronamide, L-pyroglutamic acid, and D-serine. The number of substrates shared exclusively between two species only is between one and two.

Niche overlap (NO) among all pairwise species combinations ranged from 60 to 80% (Figure 5C). *Curvibacter* sp. shares the highest overlap (80%) with *Pseudomonas* sp. and *Duganella* sp.. For the other species the overlap ranges between 60 and 70%. *Duganella* sp. displays the highest overlap with *Pseudomonas* sp. and *Undibacterium* sp. around 80%, whereas the overlap between *Pelomonas* sp. and *Acidovorax* sp. reaches almost 70%. *Undibacterium* sp. exhibits an overlap of 60 to 70% with *Acidovorax* sp., *Pseudomonas* sp. and *Pelomonas* sp.. *Acidovorax* sp. a 80% overlap with *Pseudomonas* sp. and 70% overlap with *Pelomonas* sp.. *Pseudomonas* sp. and *Pelomonas* sp. show a 75% overlap of the nutrients used. Mean niche overlap was determined as the mean of all pairwise niche overlap values for each species. Comparing the number of nutrients being used by the individual species we find that *Curvibacter* sp. and *Pseudomonas* sp. are able to use 70% of the provided substrates. *Duganella* sp. uses 57%, *Acidovorax* sp. 54% and *Pelomonas* sp. 51%. The lowest substrate utilization was measured for *Undibacterium* sp. with 41% of the available substrates.

## Discussion

Microbial communities residing in abiotic environments typically comprise numerous interacting species. Such communities have been studied with traditional approaches, for example the niche-assembly concept, which is an extension of the classical niche theory (Hutchinson, 1957). The niche-assembly perspective proposes that any ecosystem is made up of a limited number of niches, each occupied by a single species (Wennekes et al., 2012). Thus, the partitioning of these niches leads to the stable coexistence of competing species within an ecosystem. To assess the rules of assembly and coexistence of microbiota in host-associated microbiomes, we here apply the niche-assembly perspective to a metaorganism, and thus specifically extend the concept to biotic environments.

We find that the fundamental niche (here defined by the absence of interspecific microbial interactions) differs considerably from the realized niche of *Hydra*’s associated microbes (Figure 2). This reflects the difference in performance between the species when they individually occupy *Hydra* (mono-association) as to when they occur as part of their microbial community on the host. As predicted by niche theory, we find for the majority of the species that the realized niche is smaller than the fundamental one, most likely caused by interspecific microbial competition, as has also been observed in other systems, e.g. *Vibrios* in their marine environment (Materna et al., 2012). In our study, the best colonizer in the mono-colonizations, *Acidovorax* sp. (as also observed by Fraune et al. (2015)), is the least abundant species as part of the microbial community. While we cannot provide details on the nature of the interspecific competition within the whole community as observed here, a recent study investigated the interaction between the two main *Hydra* colonizers, *Curvibacter* sp. and *Duganella* sp., in more detail (Deines et al., 2020). While *Duganella* sp. likely benefitted from products excreted by *Curvibacter* sp., it was able to outcompete *Curvibacter* sp. in the tested *in vitro* environments (but not on the host). Most importantly, this effect was independent of initial frequency but depended on direct contact, which might be due to competition for the same resources or due to *Duganella* sp. actively harming *Curvibacter* sp., e.g. through contact-depending killing (see Granato et al., 2019 for an overview on potential mechanisms). In the host context, a stable co-occurrence might be achieved through spatial segregation of microbial colonizers or through active manipulation by *Hydra*, for example through the secretion of antimicrobial peptides and neuropeptides (Franzenburg et al., 2013; Augustin et al., 2017). Nevertheless, when part of the whole microbial community, both main colonizers in the community, *Curvibacter* sp. and *Duganella* sp., occupy a much bigger realized niche than fundamental niche. This finding is very interesting and indicates that the two species benefit from interactions when part of the whole microbiome. This can happen directly through positive interactions with the other members of the microbiome or indirectly by benefitting from the interactions between the other microbiome members and the host. We draw attention to the fact that the latter aspect differs from the classical Hutchinson niche concept, in that in our case the environment, i.e. the host, has the potential to change its interactions depending on the specific bacterial colonizers. Our finding also highlights the importance of the low frequency community members in shaping the overall community composition, as has recently been suggested for *Hydra* (Deines et al., 2020).

For linking the community composition in *Hydra*’s microbiome to specific characteristics, we used a trait-based approach focusing on traits potentially involved in microbiome assembly and stability (summarized in Figure 6). We draw your attention to the fact that all traits were assessed through commonly used *in vitro* assays. While this excludes interference by the host, the *in vitro* assays might not accurately reflect each bacterial trait in the host context as they might depend on and change based on the very specific external conditions that cannot be accurately mimicked with *in vitro* assays (nevertheless also note the examples given below where performance of the microbiota in the *in vitro* assays was positively associated with performance in the host).

**Figure 6.**
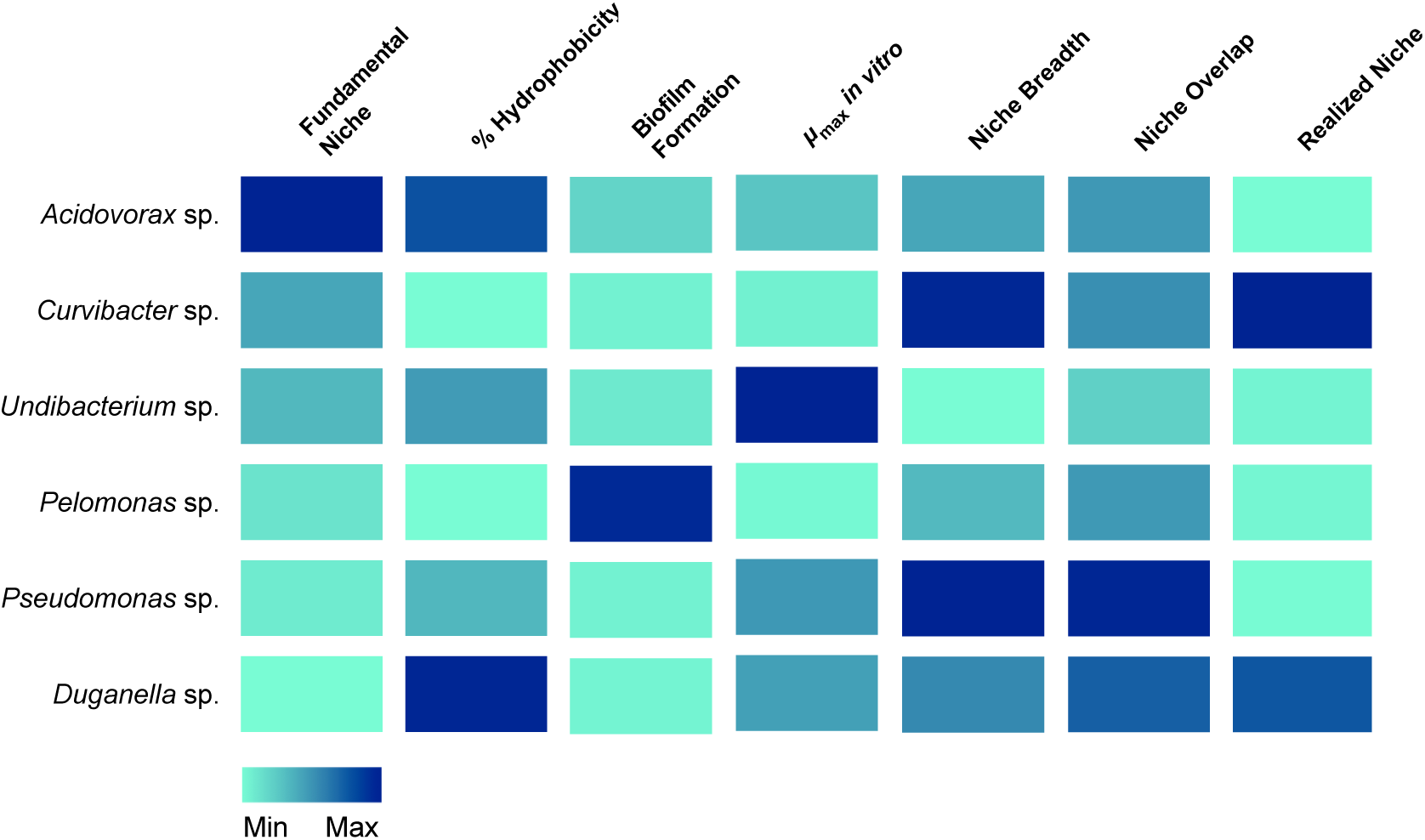
Association of traits with the occupation of the fundamental and realized niches in *Hydra*’s six microbiome members. Colors indicate the relative magnitude of the respective trait.

A first step in microbiome assembly is the attachment to host surfaces, which can happen in a multitude of ways. In the human intestine, for example, microbes have been found to bind to mucin, a major component of the human mucosa (de Vos, 2015). Adhesion is thus thought to be a powerful mechanism for exerting both, positive and negative selection for or against specific microbes (McLoughlin et al., 2016; Schluter et al., 2015). Amongst others (van Loosdrecht et al., 1987), bacterial cell surface hydrophobicity has been shown to play a crucial role in surface attachment (Krasowska and Sigler, 2014). In general, hydrophobic cells adhere more strongly to hydrophobic surfaces and vice versa (Giaouris et al., 2009; Kochkodan et al., 2008). Nevertheless, the heterogeneity of a bacterial population needs to be taken into account. For example, the presence of both, hydrophilic and hydrophobic cells, have been observed in planktonic bacteria cell populations, implying that only part of the population participates in an adhesion process to substrates (Krasowska and Sigler, 2014). We also observe mixed cell populations for most of *Hydra*’s microbial associates, except for two species, *Curvibacter* sp. and *Pelomonas* sp., which only consist of hydrophilic cells. They seem to be perfectly adapted to *Hydra*’s epithelial cells, which are coated with a carbohydrate-rich layer, the glycocalyx (Ouwerkerk et al., 2013; Schröder and Bosch, 2016). The microbiome inhabits the outer mucus-like layer of the glycocalyx (Fraune et al., 2015), which is hydrophilic. Thus, hydrophilic bacterial cells should adhere more strongly to *Hydra* than hydrophobic cells. Both, *Curvibacter* sp. and *Pelomonas* sp., have been shown to be of particular importance to the host. *Curvibacter* sp. shows signs of coevolution with its host and contributes to fungal resistance against the filamentous fungus *Fusarium* (Fraune et al., 2015). *Pelomonas* sp. has been shown to be of central importance in modulating the spontaneous body contractions in *Hydra* (Murillo-Rincón et al., 2017). So both species contribute to host fitness, providing the opportunity for the speculation that the host actively selects for specific microbes. This could happen, for example, by controlling the production and release of adhesive molecules from the host epithelium as suggested by McLoughlin et al. (2016).

After successful attachment, bacteria need to colonize the habitat. In most cases, this happens through the formation of biofilms, as has been reported for the gut (de Vos, 2015; Kania et al., 2007). The biofilm succeeds the planktonic phase in the bacterial life cycle (McDougald et al., 2012) and represents a key ecological process for the colonization of different habitats. Thus, the difference in the ability to form biofilms could provide an explanation for why one species outcompetes the other species or has a higher chance of persistence in the *Hydra* ecosystem. Further, biofilms have been shown to protect bacterial cells from various environmental stressors (Flemming and Wingender, 2010). Interestingly, from the six species tested here, the one with the highest ability to form biofilms is *Pelomonas* sp., whereas the two main colonizers, *Curvibacter* sp. and *Duganella* sp. show a reduced capacity to form biofilms. Our finding indicates that the capability of biofilm formation is not a good predictor of the bacterial performance in the *Hydra* habitat. Nevertheless, it might be of importance for the establishment and persistence of some of the low abundance species, such as *Pelomonas* sp. and *Acidovorax* sp..

Importantly, microbiomes on external surfaces of metaorganisms, such as the skin, have been reported to be highly stable despite their constant exposure to extrinsic factors (Oh et al., 2016). Whereas bacterial diversity is widely recognized in leading to temporal stability of ecosystem processes (Bell et al., 2009; Griffin et al., 2009; Prosser et al., 2007), the influence of resource niche breadth has received little scientific attention (Hunting et al., 2015). Recent work studying the decomposition of organic matter in experimental microcosms found that the higher the overlap in resource niches, the higher the stability of the microbial community. It is reasonable to assume that the same underlying principles govern stability in host-associated microbial communities. We therefore measured the niche overlap and resource use of the six species isolated from the *Hydra* microbiome. Interestingly, we find the niche overlap between all pairwise combinations to be between 60 and 80%, with about 20% of the carbon sources being metabolized by all species. This suggests that metabolic overlap could be involved in promoting the extreme temporal stability of *Hydra*’s microbiome (Fraune and Bosch, 2007) in addition to active manipulation by *Hydra* through the secretion of antimicrobial peptides and neuropeptides (Franzenburg et al., 2013; Augustin et al., 2017). We also found the two main colonizers, *Curvibacter* sp. and *Duganella* sp. together with *Pseudomonas* sp., to possess the widest resource niche breadth of all species, and that five out of six species were able to metabolize more than 50% of the 95 offered carbon substrates. Overall, the relative niche breadth observed in the tested species can serve as a proxy of the metabolic diversity of the *Hydra* microbiome. In the metaorganism *Hydra* it seems that niche breadth of its symbionts is a fairly good indicator for relative microbial performance as compared to the other members of the community, and thus of the realized niche (Figure 6). This is not the case for *Pseudomonas* sp., which despite its observed niche breadth, does not seem to play a major role as part of *Hydra*’s microbiome. One reason for this might stem from the fact that *Pseudomonas* sp. is an ubiquitous bacterium, which is generally characterized by its ability to colonize all major environments and might well be the bacterial species with the broadest ecological niche range (Spiers et al., 2000).

The metabolic overlap, i.e. redundancy, within *Hydra*’s microbial community indicates that the individual species are not occupying a specific metabolic niche. Nevertheless, the only one for which we observed a specific carbon usage pattern is the main colonizer *Curvibacter* sp., which utilizes eight carbon sources that are not metabolized by any other tested microbiome members. Whether this hints at the occupation of a specific niche within *Hydra*’s microbial community and can be linked to the observation that its realized is bigger than its fundamental niche is currently open to speculation. An alternative option might be that *Curvibacter* sp. is auxotrophic in producing certain amino acids (four of the uniquely used carbon sources are the amino acids L-Leucine, L-Phenylalanine, L-Pyroglutamic Acid, and D-Serine), as are 98% of all sequenced microbes (Zengler and Zaramela, 2018). *Curvibacter* sp. might thus rely on the uptake of external substrates that might not be secreted by the host but by its fellow community members. Analyzing the metabolic interactions within this microbial network will be essential for understanding community assembly, composition, and maintenance.

In summary we find that the here measured bacterial traits vary across microbiome members. Further, the dominant species in the microbiome do not necessarily perform best in all of the measured traits. We rather observe that all species, independent of their density, perform well in a subset of traits, likely facilitating the coexistence of several niches within the host ecosystem. Whether a change in the realized niche of microbes can be linked to potential for dysbiosis is an interesting aspect, which warrants further investigation.

## Ethics statement

Ethical restrictions do not apply to cnidarian model organisms such as *Hydra*.

## Author contributions

PD and KH designed the experiments. PD performed the experiments. PD and KH analysed the data. PD, KH and TB wrote the paper.

## Acknowledgements

TB gratefully appreciates support from the Canadian Institute for Advanced Research (CIFAR) and thanks the Wissenschaftskolleg (Institute of Advanced Studies) in Berlin for a sabbatical leave.

